# Independent domestications shape the genetic pattern of a reproductive isolation system in rice

**DOI:** 10.1101/2020.01.08.898130

**Authors:** Xun Xu, Song Ge, Fu-Min Zhang

## Abstract

Severe reproductive isolation (RI) exists between the two subspecies of rice, *Indica* and *Japonica*, but in the wild ancestors no post-zygotic RI was found. The studies about the establishment of the interspecies RI of rice are still rear. A pair of rice hybrid sterility genes, *DOPPELGANGER 1* (*DPL1*) and *DOPPELGANGER 2* (*DPL2*), offers a convenient example to study the evolutionary history of RI genes. Either of the two loci has one non-functional allele (*DPL1-* and *DPL2-*). The hybrid pollen carrying both *DPL1-* and *DPL2-* will be sterility.

We collected 811 individuals: *Oryza sativa* (132), the two wild ancestors *O. nivara* (296) and *O. rufipogon* (383) as well as 20 *DPL1* and 34 *DPL2* sequences of *O. sativa* from on-line databases. We analysed the genetic and geographic pattern of *DPLs* in all three species to determine the origination regions of *DPL1-* and *DPL2-*. The neutral test as well as the diversities of nucleotide and haplotype were used to detect if selection shaped the pattern of *DPLs*.

We found that *DPL1-* and *DPL2-* of rice emerged from wild ancestor populations in South Asia and South China through two respective domestications. Comparing with the ancestral populations, *DPL1-* and *DPL2-* both showed reduce of diversities, however their frequencies increased in rice. We assume that the reduce of diversities due to the bottleneck effect of domestication while the loss of one copy was preferred by artificial selection for cost savings.

## Introduction

Reproductive isolation (RI) is considered a failure to transmit genetic materials (Futuyma, 2013). Barriers that lead to RI are classified as pre- or postzygotic for practical (Baack et al, 2015; Futuyma, 2013; Lafon-Placette et al, 2016; Rieseberg and Willis, 2007; Widmer et al, 2009). Many genes related to RI have been identified in various species (Fishman and Sweigart, 2018; Masly et al, 2006; Nadir et al, 2018; Ouyang and Zhang, 2018; Zuellig and Sweigart, 2018), yet studies that aim to elucidate the evolutionary histories of RI genes are still rare.

Rice (*Oryza sativa*) is an important crop as well as a model species. Strong postzygotic RI was found between two subspecies of rice, *Japonica* and *Indica*. Many loci answering for the hybrid incompatibility have been found (Ouyang et al, 2010; Wang et al, 2014). Hundreds of loci were found related to hybrid incompatibility, indicating that the incompatibility between *Japonica* and *Indica* has a complex genetic background (Li et al, 2017). Until now, three independent system were detected: *S5* (Chen et al, 2008; Yang et al, 2012b), *Sa* (Long et al, 2008) and *DPLs* (Mizuta et al, 2010). *S5* leads to endoplasmic reticulum stress. *Sa* and *DPLs* both lead to hybrid pollen sterility. On the contrary, the two wild ancestors of rice, *O. nivara* and *O. rufipogon*, are compatible (Cai et al, 2019). Rice and the two wild ancestors constitute a excellent system for studying the evolution history of RI genes.

The *DPL* system offers a suitable example for exploring this phenomenon. The *DPL* system contains only two genes, *DOPPELGANGER 1* (*DPL1*) and *DOPPELGANGER 2* (*DPL2*), causes pollen sterility by reciprocal gene loss (Mizuta et al, 2010). We consider it as a tidy example for thorough evolutionary study. Both genes of *DPL* system have two alleles: *Indica-*like *DPL1* (non-functional) and *Japonica-*like *DPL1* (functional), *Indica-*like *DPL2* (functional) and *Japonica-*like *DPL2* (non-functional). Hybrids carrying non-functional alleles at both loci exhibit semi-sterility of pollen. For simplicity, we classified the alleles as functional (*DPL1+*/*DPL2+*) and non-functional (*DPL1-*/*DPL2-*) according to Mizuta et al (2010).

The genetic pattern of *DPLs* in *O. sativa* remains unknown, let alone its wild ancestors. Rice includes 6 groups (Wang et al, 2014): *Japonica* can be divided into *temperate japonica*, *tropical japonica*, *aromatic* and *rayada*, while *Indica* consists of *Indica* and *aus*. Previous studies indicated that *DPL1-* was fixed in *aus* but relatively rare in *indica*, while *DPL2-* was common in the groups of *Japonica* (Craig et al, 2014; Mizuta et al, 2010). Among the wild rices, *DPL1-* was detected but *DPL2-* was not. The absence of *DPL2-* in wild rices indicates that *DPLs* were established during domestication, not inherited. Nevertheless, both studies are short of population sampling. In addition, *aromatic* and *rayada* are minor groups of *Japonica* and were ignored in previous studies.

In our study, we take *DPLs* as an example to release the establishment of RI between *Indica* and *Japonica*. To be specific, we ask 1) Do the *DPL* system exists in wild rice? 2) If so, how can they are maintained since hybrid sterility leads to the waste of reproductive resources? 3) How the *DPL* system originated and what is the role of domestication?

## Materials and Methods

### Plant materials

We sampled 296 *O. nivara* and 383 *O. rufipogon*. The whole natural distribution areas of wild ancestors were covered (Fig. 1). The materials are listed in Additional file 1.

**Fig. 1.**
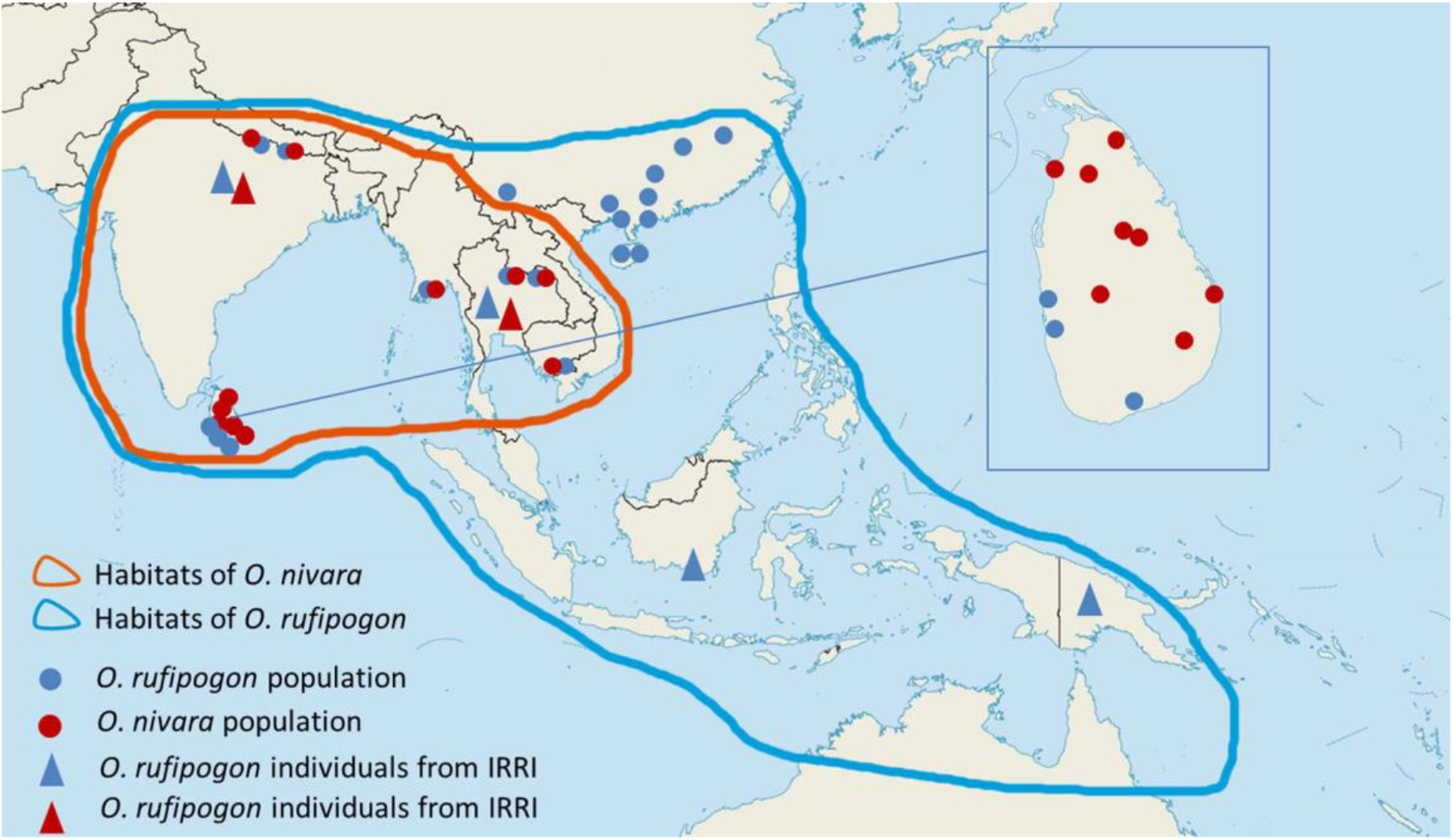
The distribution of wild rice samplings, functional and non-functional alleles of *DPLs*. The populations of *O. nivara* and *O. rufipogon* are respected with spots. Triangle roughly shows the origin of individuals from IRRI. Red and blue lines indicate the natural habits of *O. nivara* and *O. rufipogon*, respectively. Circles indicate frequencies of functional and non-functional alleles.

Since South Asia is closely related with the domestication of rice (Garris et al, 2005; Huang et al, 2012; Morishima et al, 1992), ten populations have been collected from South Asia and Southeast Asia, including Nepal (NEP), Myanmar (MMR), Cambodia (KHM) and Laos (LAO). The number of individuals in the populations ranges from 29 to 36.

We also collected *O. rufipogon* from New Guinea, which is an distinct line and is named Oceanian type (Oka, 1988). we further included a total of 115 individuals from the International Rice Research Institute (IRRI), including 6 populations from India (IND), Thailand (THA), Indonesia (IDN) and New Guinea (PNG). To increase representation, eleven populations were collected from Sri Lanka (LKA), each including 9-10 individuals. Ten Chinese *O. rufipogon* populations were contained, each including 10-11 individuals. One population was from Jiangxi (JXDX) province of China, the northernmost frontier of natural habitats of *O. rufipogon* (Morishima et al, 1992). Two populations were from Hunan (HNCL, HNJY), the province near Jiangxi. One population from Guangdong (GDGZ) and three populations from Guangxi (GXHZ, GXBH, GXTD) were collected from South China, where are related with the domestication of *Japonica* (Sang and Ge, 2007). Two populations were from Hainan (HNDZ, HNWC), a tropical island in the natural habitat of *O. rufipogon*. One population from Yunnan (YNJH), where are the natural habitats of *O. rufipogon* and several other *Oryza* species. In addition, 131 *O. sativa* landraces were included from all 6 groups: *aromatic* (ARO), *aus* (AUS), *indica* (IND), *rayada* (RAY), *temperate japonica* (TEJ) and *tropical japonica* (TRJ).

### Sequencing and genotype analysis

We extracted genomic DNA from fresh or silica gel-desiccated leaves with a DNAsecure Plant Kit (TIANGEN, Beijing, China). The primers and thermal cycling procedure were as described by Craig et al (2014). The genomic DNA and PCR products were tested with gel electrophoresis. PCR products were sanger sequenced (Majorbio, Shanghai, China). When any dual peaks occurred, indicating heterozygous individuals, the PCR products were cloned with pEASY-T1 vectors (TransGen Biotech, Beijing, China). We consulted previous studies to confirm singletons (Zheng and Ge, 2010). All sequences were submitted to GenBank. We downloaded 20 *DPL1* and 34 *DPL2* sequences for *O. sativa* from the National Center for Biotechnology Information (NCBI, https://www.ncbi.nlm.nih.gov) (Additional file 2: Table S2).

For simplicity, *DPL1-* and *DPL2-* were defined according to previous study (Mizuta et al, 2010). *DPL1-* was defined by a 518 bp transposable element insertion in the first coding sequence (CDS), while *DPL2-* was defined by a single nucleotide transition from "A" to "G" in the intron, leading to a premature stop codon and producing a readthrough protein (Additional file 3: Figure S1).

### Genetic diversity and phylogeographic analysis

All obtained sequences were aligned with Clustal X 1.83 (Thompson et al, 1997) and revised manually with BioEdit 7.0.9.0 (Hall, 1999). To detect potential trace of selection, we evaluated the number of polymorphic sites (S), nucleotide diversity (π) (Nei, 1987) and performed neutrality tests (Tajima’s D (Tajima, 1989)) with DNASP 5.1 (Librado and Rozas, 2009). The number of haplotypes (h) and haplotype diversity (Hd) were also calculated with DNASP 5.1 after excluding recombined sequences with the online tool IMgc (Woerner et al, 2007). Insertions and deletions were excluded during the analysis of diversities. The difference of nucleotide diversity between each groups of *O. sativa* and wild ancestors was performed as the relative ratio of π in each groups to π in wild ancestor populations. The difference of haplotype diversity (Hd) was performed in the same way.

We draw network to verify the exact emergence times and origination centre of *DPL1* and *DPL*2 in rice. Median-joining method (Bandelt et al, 1999) was used to construct networks of haplotype with Network 5011 (Fluxus Technology Ltd.). Recombined sequences were excluded with IMgc (Woerner et al, 2007) to clarify the skeleton. MP Calculation (Polzin and Daneshmand, 2003) was used to simplify the rare haplotypes.

## Results

### Allelic Distribution

A total of 695 *DPL1* and 787 *DPL2* sequences were obtained. In addition, we collected 20 *DPL1* sequences and 34 *DPL2* sequences from online databases (Craig et al, 2014; Mizuta et al, 2010) (Additional file 2: Table S2). The alignments with gaps were 1407 bp and 439 bp in length, respectively. All sequences are available in GenBank (http://www.ncbi.nlm.nih.gov) under the accessions MN446022 - MN446138, MN446139 - MN446234, MN446235 - MN446531, MN446532 - MN446716, MN476103 - MN476405, MN476406 - MN476830, MN476831 - MN476868, MN476869 - MN476888.

The frequencies of *DPL1-* and *DPL2-* are in Table 1. *DPL1-* in *O. sativa* (10.2%) is in between of the two ancestors, *O. nivara* (37%) and *O. rufipogon* (5.0%). *DPL2-* existed only in one wild rice (*O. rufipogon*) with a frequency of 4.9%, while in *O. sativa* the frequency increased to 32.3%. In contrast to other studies (Craig et al, 2014; Mizuta et al, 2010), we found both *DPL1-* and *DPL2-* in wild rices, indicating that the *DPL* system already existed before the domestication of rice.

**Table 1.** Nucleotide polymorphism and neutral test of *O. rufipogon*, *O. nivara* and *O. sativa*.

| Population/<br>Species | Number | | Frequency | | S | | $\pi$ | | h | | Hd | | Tajima's D | |
| --- | --- | --- | --- | --- | --- | --- | --- | --- | --- | --- | --- | --- | --- | --- |
|  | DPL1- | DPL2- | DPL1- | DPL2- | DPL1 | DPL2 | DPL1 | DPL2 | DPL1 | DPL2 | DPL1 | DPL2 | DPL1 | DPL2 |
| *O. nivara* | 91 | 0 | 37.0% | 0.0% | 41 | 12 | 0.00336 | 0.0009 | 23 | 9 | 0.819 | 0.267 | -1.73831 | **-1.8546** |
| *O. rufipogon* | 16 | 18 | 5.0% | 4.9% | 79 | 34 | 0.00421 | 0.0029 | 53 | 25 | 0.914 | 0.429 | **-2.1935** | **-2.11384** |
| *O. sativa* | 15 | 49 | 10.2% | 30.4% | 11 | 6 | 0.00118 | 0.00363 | 7 | 4 | 0.223 | 0.571 | -1.21896 | 1.04607 |
| nIND | 18 | 0 | 45.0% | 0.0% | 12 | 5 | 0.0021 | 0.0014 | 6 | 5 | 0.638 | 0.41 | -1.18432 | -1.17921 |
| rIND | 2 | 0 | 11.8% | 0.0% | 17 | 8 | 0.00422 | 0.00292 | 9 | 7 | 0.955 | 0.584 | -1.31101 | -1.22445 |
| nTHA | 5 | 0 | 62.5% | 0.0% | 6 | 3 | 0.0037 | 0.00105 | 3 | 3 | 0.762 | 0.295 | 1.59761 | -1.65231 |
| rTHA | 2 | 0 | 13.3% | 0.0% | 12 | 3 | 0.00345 | 0.0011 | 5 | 3 | 0.813 | 0.362 | -1.08464 | -1.34917 |
| nKHM | 29 | 0 | 93.5% | 0.0% | 11 | 1 | 0.00202 | 0.00014 | 5 | 2 | 0.743 | 0.125 | -1.19393 | -1.14244 |
| rKHM | 6 | 0 | 22.2% | 0.0% | 6 | 0 | 0.00182 | 0 | 5 | 4 | 0.667 | 0.2 | -0.48253 | NA |
| nMMR | 0 | 0 | 0.0% | 0.0% | 0 | 3 | 0 | 0.0005 | 1 | 2 | 0 | 0.063 | NA | -1.56135 |
| rMMR | 0 | 0 | 0.0% | 0.0% | 22 | 10 | 0.00429 | 0.00361 | 9 | 1 | 0.714 | 0 | -1.52558 | -1.00632 |
| nNEP | 0 | 0 | 0.0% | 0.0% | 4 | 1 | 0.00083 | 0.00028 | 3 | 3 | 0.53 | 0.17 | -0.76705 | -0.77374 |
| rNEP | 2 | 0 | 7.1% | 0.0% | 5 | 3 | 0.00123 | 0.00071 | 3 | 7 | 0.574 | 0.696 | -0.51862 | -1.37016 |
| nLAO1 | 20 | 0 | 100.0% | 0.0% | 13 | 1 | 0.00341 | 0.00015 | 4 | 2 | 0.591 | 0.067 | 1.06173 | -1.147 |
| rLAO1 | 0 | 0 | 0.0% | 0.0% | 12 | 13 | 0.00278 | 0.0024 | 8 | 5 | 0.837 | 0.261 | -1.04354 | **-2.24274** |
| nLAO2 | 19 | 0 | 90.5% | 0.0% | 8 | 0 | 0.00142 | 0 | 4 | 1 | 0.348 | 0 | -1.54473 | NA |
| rLAO2 | 1 | 0 | 3.4% | 0.0% | 13 | 2 | 0.00297 | 0.00102 | 7 | 2 | 0.817 | 0.228 | -0.78196 | -0.24788 |
| nLKA04 | 0 | 0 | 0.0% | 0.0% | 1 | 0 | 0.00023 | 0 | 2 | 1 | 0.2 | 0 | -1.11173 | NA |
| nLKA05 | 0 | 0 | 0.0% | 0.0% | 3 | 0 | 0.00187 | 0 | 2 | 1 | 0.533 | 0 | 1.83053 | NA |
| nLKA07 | 0 | 0 | 0.0% | 0.0% | 2 | 0 | 0.00047 | 0 | 3 | 1 | 0.378 | 0 | -1.40085 | NA |
| nLKA08 | 0 | 0 | 0.0% | 0.0% | 1 | 0 | 0.00026 | 0 | 2 | 1 | 0.222 | 0 | -1.8823 | NA |
| nLKA09 | 0 | 0 | 0.0% | 0.0% | 0 | 0 | 0 | 0 | 1 | 1 | 0 | 0 | NA | NA |
| nLKA10 | 0 | 0 | 0.0% | 0.0% | 0 | 3 | 0 | 0.00198 | 1 | 3 | 0 | 0.6 | NA | -0.65748 |
|  | DPL1- | DPL2- | DPL1- | DPL2- | DPL1 | DPL2 | DPL1 | DPL2 | DPL1 | DPL2 | DPL1 | DPL2 | DPL1 | DPL2 |
| nLKA11 | 0 | 0 | 0.0% | 0.0% | 3 | 0 | 0.00098 | 0 | 3 | 1 | 0.417 | 0 | -0.93613 | NA |
| nLKA12 | 0 | 0 | 0.0% | 0.0% | 4 | 2 | 0.00201 | 0.00091 | 2 | 1 | 0.429 | 0 | 0.48523 | -1.40085 |
| rLKA01 | 0 | 0 | 0.0% | 0.0% | 3 | 1 | 0.00195 | 0.00076 | 4 | 2 | 1 | 0.333 | 0.16766 | -0.93302 |
| rLKA06 | 0 | 0 | 0.0% | 0.0% | 1 | 0 | 0.00033 | 0 | 1 | 1 | 0 | 0 | -1.00623 | NA |
| rLKA13 | 0 | 0 | 0.0% | 0.0% | 0 | 1 | 0 | 0.00046 | 1 | 1 | 0 | 0 | NA | -1.11173 |
| rGDGZ | 0 | 10 | 0.0% | 47.6% | 8 | 4 | 0.00152 | 0.00354 | 7 | 3 | 0.569 | 0.648 | -1.48037 | 1.12924 |
| rGXBH | 0 | 0 | 0.0% | 0.0% | 6 | 2 | 0.00285 | 0.00198 | 2 | 2 | 1 | 0.4 | -0.49605 | -0.05002 |
| rGXHZ | 0 | 8 | 0.0% | 34.8% | 8 | 4 | 0.00213 | 0.00236 | 6 | 4 | 0.748 | 0.679 | -0.6049 | -0.1232 |
| rGXTD | 0 | 0 | 0.0% | 0.0% | 18 | 2 | 0.0044 | 0.0023 | 11 | 2 | 0.912 | 0.5 | -1.13215 | 1.81115 |
| rHNCL | 0 | 0 | 0.0% | 0.0% | 7 | 2 | 0.00235 | 0.00249 | 4 | 3 | 0.81 | 0.607 | -0.8501 | 1.82766 |
| rHNJY | 0 | 0 | 0.0% | 0.0% | 11 | 3 | 0.00407 | 0.00294 | 3 | 3 | 0.6 | 0.679 | -0.38658 | 0.77501 |
| rHNDZ | 0 | 0 | 0.0% | 0.0% | 8 | 4 | 0.00306 | 0.0037 | 2 | 3 | 0.476 | 0.733 | -0.51253 | 0.56555 |
| rHNWC | 0 | 0 | 0.0% | 0.0% | 11 | 5 | 0.00332 | 0.00332 | 7 | 4 | 0.944 | 0.75 | -0.23157 | -0.56682 |
| rJXDX | 0 | 0 | 0.0% | 0.0% | 8 | 2 | 0.00331 | 0.00213 | 6 | 2 | 0.952 | 0.533 | -0.25553 | 1.03299 |
| rYNJH | 0 | 0 | 0.0% | 0.0% | 5 | 1 | 0.00251 | 0.00046 | 5 | 2 | 0.806 | 0.2 | 0.28702 | -1.11173 |
| rIDN | 0 | 0 | 0.0% | 0.0% | 5 | 4 | 0.00304 | 0.00365 | 4 | 3 | 1 | 0.7 | 0.56199 | -1.0938 |
| rPNG | 3 | 0 | 60.0% | 0.0% | 5 | 2 | 0.00281 | 0.00178 | 3 | 2 | 0.7 | 0.4 | 0 | 0.1959 |
| sARO | 0 | 1 | 0.0% | 25.0% | 0 | 4 | 0 | 0.00456 | 1 | 2 | 0 | 0.5 | NA | -0.78012 |
| sAUS | 12 | 1 | 70.6% | 4.8% | 7 | 3 | 0.00286 | 0.00068 | 4 | 2 | 0.596 | 0.1 | 0.61839 | -1.72331 |
| sIND | 2 | 5 | 3.0% | 7.5% | 5 | 5 | 0.00035 | 0.00135 | 2 | 3 | 0.06 | 0.252 | -1.64155 | -0.99668 |
| sRAY | 0 | 2 | 0.0% | 66.7% | 1 | 3 | 0.00078 | 0.00456 | 2 | 2 | 0.667 | 0.667 | NA | NA |
| sTEJ | 0 | 36 | 0.0% | 87.8% | 1 | 4 | 0.00007 | 0.00172 | 2 | 3 | 0.056 | 0.292 | -1.13321 | -0.46012 |
| sTRJ | 1 | 4 | 4.8% | 16.0% | 7 | 5 | 0.00078 | 0.00417 | 3 | 4 | 0.186 | 0.684 | **-2.13123** | 1.11236 |

*DPL1-* and *DPL2-* exhibited clear structure among groups of rice. Only *aus*, *indica* and *tropical japonica* carried *DPL1-*, with frequencies of 70.6%, 3% and 4.8%, respectively. *DPL2-* was detected in all 6 groups of *O. sativa*, with the highest frequency in *temperate japonica* (87.8%) and the lowest frequency in *aus* rice (4.8%). We assume that *DPL* system is more likely a RI system between *aus* and *temperate japonica*, rather than between the two subspecies.

### Phylogeography

The distribution of non-functional alleles among populations confirmed that strong geographic structure of *DPL1* and *DPL2* (Table 1). *DPL1-* was found in both *O. nivara* and *O. rufipogon* but clustered mostly in South Asia (India and Nepal) and Southeast Asia (Thailand, Cambodia and Laos). *DPL2-* only existed in two *O. rufipogon* populations in southern China (rGDGZ and rGXHZ). With geographic isolation, the incompatible alleles failed to trigger hybrid fertility in natural condition. Hence the *DPL* system of RI can maintain for long in wild rices.

The network of *DPL1* proved that the *DPL1-* of *O. sativa* clustered only in H_1 (Fig. 2a). The result indicates a once-emergence from the wild ancestors containing H_1, on the contrary of the multiple emergence hypothesis from previous study (Mizuta et al, 2010). The two most frequent haplotypes of *DPL1+* of rice (H_5 and H_36) were immediately related with each other but separated from H_1, proves that the originations of *DPL1-* and *DPL1+* were independent.

**Fig. 2.**
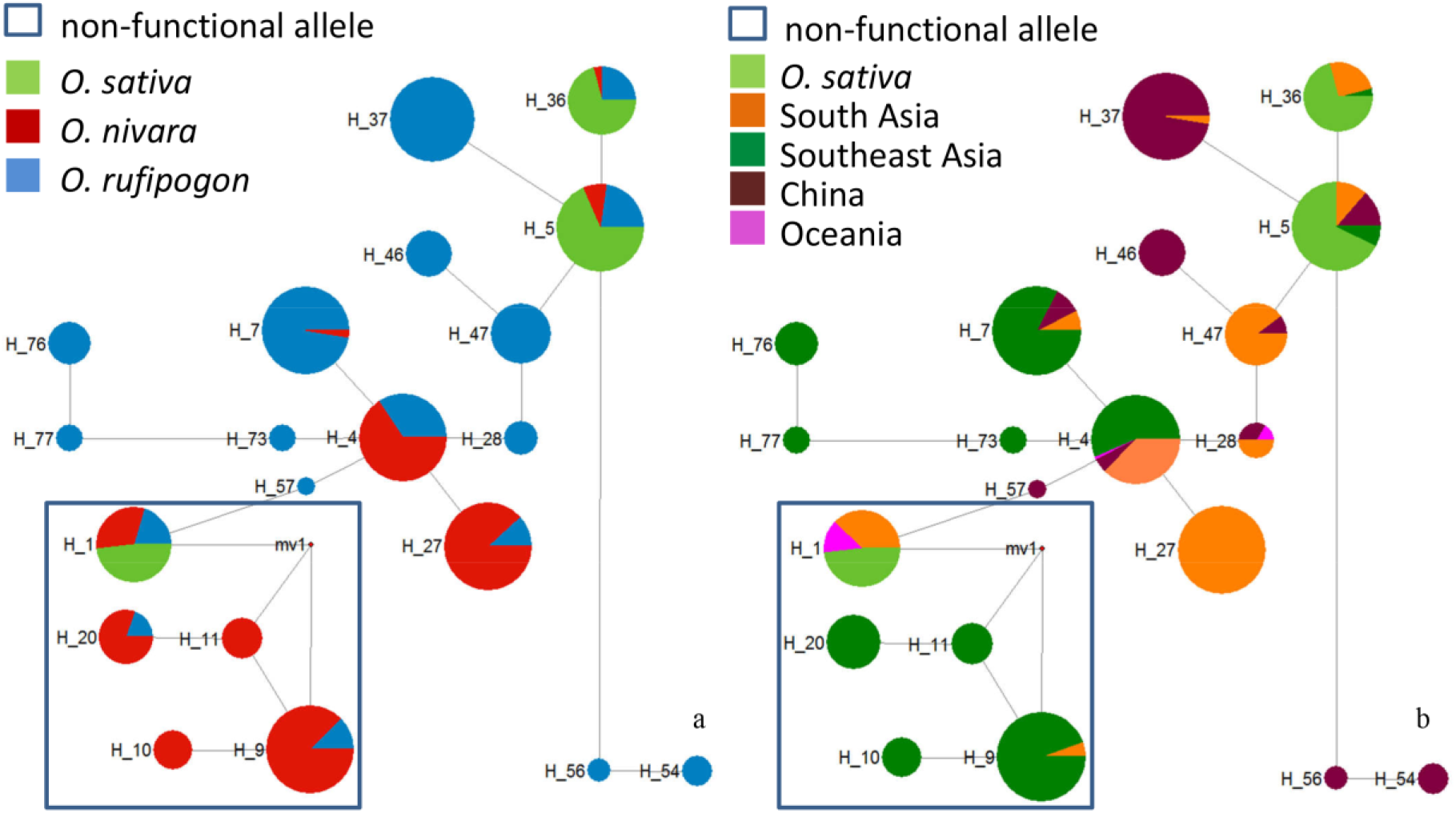
The median-joining haplotype networks of *DPL1*. Sections of each circle represent the proportion species (a) or locations (b) possessing in each haplotype. Black dots represent possible missing haplotypes. Size of each circle represent the number of sequences of haplotype. The length of grey lines bases on the number of mutation. Non-functional haplotypes are emphasized with hollow-square.

Fig. 3a indicates that all *DPL2-* sequences of rice were concentrated in H_13 and that the *DPL2+* of rice appeared in H_1 and H_5. Similar with *DPL1*, the *DPL2+* and *DPL2-* of rice were separated. We consider that the *DPL2-* and *DPL2*+ of rice also originated respectively.

**Fig. 3.**
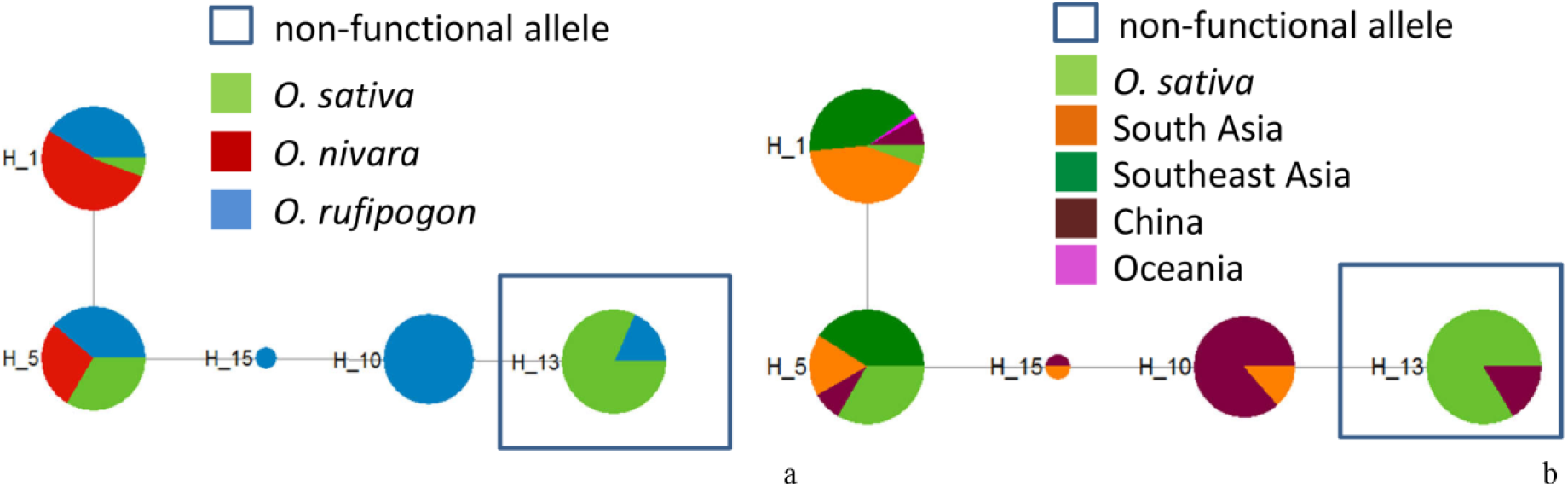
The median-joining haplotype networks of *DPL2*. Sections of each circle represent the proportion species (a) or locations (b) possessing in each haplotype. Black dots represent possible missing haplotypes. Size of each circle represent the number of sequences of haplotype. The length of grey lines bases on the number of mutation. Non-functional haplotypes are emphasized with hollow-square.

According to Fig. 2b, the *DPL1-* in wild rices shows discontinuous geographic distribution. H_1 contains mostly the *O. nivara* from India (nIND) and *O. rufipogon* from Oceania (rPNG). Previous studies proved that South Asia is a domestication region of rice (Kovach et al, 2007; Sweeney and McCouch, 2007; Vaughan et al, 2008). Therefore, we assume the *DPL1-* of *O. sativa* is from nIND. H_5 and H_36 cover the wild rices from almost every region, indicating that *DPL1+* occurred before *O. sativa* and the two wild species diverged.

H_13 includes haplotypes from two ajoining *O. rufipogon* populations: rGDGX and rGXHZ (Fig. 3b). H_1 and H_5 covered samples from every habitats of wild ancestors, indicating that *DPL2+* exists in wild rices for a long time. Therefore, we assume that *DPL2-* originated from rGDGX and rGXHZ then was inherited by rice.

In conclusion, *DPL1-* and *DPL2-* of *O. sativa* are both immediate successors from wild ancestors, originated at geographically isolated areas: *DPL1-* originated from *O. nivara* in India while *DPL2-* originated from *O. rufipogon* in South China. The origination regions of *DPL1+* and *DPL2*+ is still uncertain because of their wide distributions.

### Tests for selection

#### Frequency variance of *DPL1-* and *DPL2-*

Since the two wild rice populations containing *DPL2-* (rGDGZ and rGXHZ) adjoined, we emerged them into one ancestor population rGDGX. We compared the frequencies of *DPL1-* and *DPL2-* with their ancestor populations (Table 1). The frequency of *DPL1-* of *aus* is 70.6%, higher than nIND (45.0%). The frequency of *DPL2-* of *temperate japonica* is 87.8%, also higher than rGDGX (40.9%). We assume that *DPL1-* and *DPL2-* were preferred during artificial selection and the frequencies increased as a consequence.

#### Neutral test

The effect of domestication comes from neutral bottleneck and artificial selection. We used Tajima’s D to test if *DPL1* and/or *DPL2* underwent artificial selection, and if the selection differed among populations/groups (Table 1). In *O. rufipogon*, purifying selection was detected in both *DPL1* (−2.1935, p<0.01) and *DPL2* (−2.11384, p<0.01). Only *DPL2* was under purifying selection in *O. nivara* (−1.8546, p<0.05). Both loci were under neutral condition in *O. sativa*. However, at the population level, we only found significant purifying selection on *DPL2* in rLAO1 (−2.24274, p<0.05). Both genes were under neutral selection in the other populations. Although not significant, the values of Tajima’s D of most populations/groups are negative. Besides, We found the lack of polymorphic sites (S) in several populations/groups (Table 1). The purifying selection was a possible reason.

According to the complex pattern among populations, it is hasty to draw a conclusion that the significant results of selection at species level were caused by selection. One of the basic hypothesis of neutral tests is that the sample must be a population with internal gene flow (Tajima, 1989). However, the two wild species include many geographical isolated populations. We assume that the significance Tajima’s D at species level may be caused by intraspecies structure or population-specific rare haplotypes.

#### Nucleotide and haplotype diversity analysis

We then estimated the nucleotide diversity (π) of *DPL1* and *DPL2* in all three species and each population/group (Table 1). In the two wild rices, we found the nucleotide diversity of *DPL1* was higher than that of *DPL2* (*O. nivara*: 0.00336>0.0009; *O. rufipogon*: 0.00421>0.0029). In *O. sativa* we found the opposite: the nucleotide diversity of *DPL1* (0.00118) was lower than that of *DPL2* (0.00363). After excluding recombined sequences, the number of recombined sequence, haplotype diversity (Hd) are listed in Table 1. The Hd of *DPL1* in *O. sativa* is 0.223, lower than *DPL*2 (0.571). In *O. nivara* and *O. rufipogon* the Hd of *DPL1* are higher then *DPL*2: 0.819>0.267; 0.914>0.429. The patterns of π and Hd also confirm the affect of domestication.

We then compared the nucleotide and haplotype diversities of *DPL1* between *aus* and wild ancestral population nIND, then compared those of *DPL1-*. The same comparison was used on *DPL*2 and *DPL2-* between *temperate japonica* and rGDGX.

The results are in Fig. 4.

**Fig. 4.**
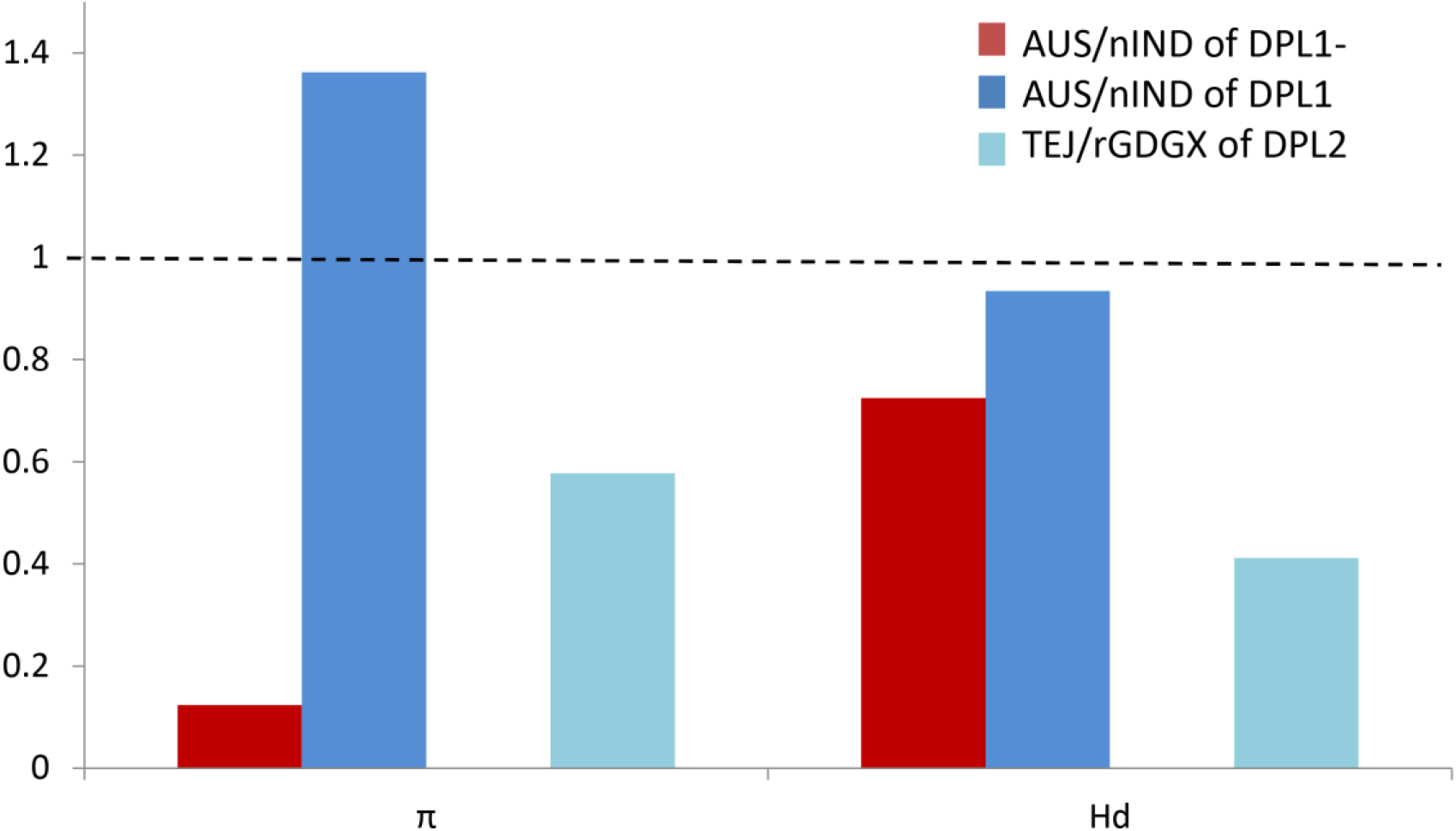
The ratio of nucleotide diversity (π) and haplotype diversity (Hd) between groups of *O. sativa* and the ancestral populations. AUS and TEJ indicate aus rice and temperate japonica, respectively. The nIND and rGDGX indicate the two ancestral populations: *O. nivara* from India and the two *O. rufipogon* populations carrying *DPL2-* from South China.

We unexpectedly found the *DPL1* of *aus* contained more nucleotide diversity (136.19%) than nIND. The haplotype diversity of *aus* (0.596) and nIND (0.638) were almost equal and the ratio is 93.42%. The *DPL1-* of *aus* contained 12.37% of nucleotide diversity but the number of haplotypes (3) was equal to nIND and the ratio of haplotype diversity was 72.43%. We found the *DPL*2 of *temperate japonica* contained 57.71% of nucleotide diversity and 41.18% of haplotype diversity of rGDGX. The *DPL2-* of *temperate japonica* showed severer reduce of diversity, contained only one haplotype and no diversity (indicated with space in Fig. 4).

In conclusion, we assume the haplotypes of *DPL1-* and *DPL2-* in *O. sativa* were randomly reserved by bottleneck, then the artificial selection lead to the increase of frequencies. Besides, *DPL*2 underwent severer pressure during domestication than *DPL1*.

## Discussion

### Domestication shaped *DPL* system from wild ancestors

The RI between *Japonica* and *Indica* has been widely reported (Morishima et al, 1992; Ouyang and Zhang, 2013), but the ancestors, *O. nivara* and *O. rufipogon* were proved compatible (Cai et al, 2019). Here comes the question that whether the RI appeared during the process of domestication, or the potential RI already existed in the ancestors.

Until now, three major system were detected: *S5* (Chen et al, 2008; Yang et al, 2012b), *DPLs* (Mizuta et al, 2010) and *Sa* (Long et al, 2008). The *S5* system can be roughly classified into three types: *Indica-*like, *Japonica-*like and wide compatibility. *Indica-*like and *Japonica-*like are incompatible, and wide compatibility shows compatible with both of the others. All three types exist in wild rice (Craig et al, 2014; Du et al, 2011), which provides an evidence of potential RI. *Sa* causes hybrid pollen sterility, includes two adjacent genes *SaM* and *SaF*. *Indica* and *Japonica* carries *SaM+SaF+* and *SaM-SaF-*, respectively. *SaM+*, *SaF+* and *SaM-* are all necessary for pollen sterility, and discovered in wild rices (Craig et al, 2014; Long et al, 2008). These studies indicate that some potential reproductive barriers between *Japonica* and *Indica* have existed in their ancestors, yet rise another questions that how the incompatible alleles maintaining natural condition. The next question is how domestication shapes the potential materials. Ouyang and Zhang (2018) suggested three models to explain the establishment of the incompatibility within rice. However, the models focused on how the incompatible loci accumulated in each divergent linage, and may not be accurate for some locus due to the lack of clear evolutionary background information. For example, Ouyang and Zhang (2018) thought that *DPL2-* did not exist in wild rices and believed that the incompatibility arose during the divergence of the two subspecies.

Our study for the first time discovered both *DPL1-* and *DPL2-* in wild rice. Therefore, the three major systems are all heritages rather than de novo mutations during the domestication.

We hence hypothesize that the RI between *Indica* and *Japonica* may contain a more complex genetic background than what we have known. Researches on larger scale and involving more groups of rice are helpful to understand the whole scene. With one typical *Japonica* and two varietal groups of *Indica*, 43 loci and 223 interactions were located, which are involved in the fertility of embryo-sac, pollen and spikelet (Li et al, 2017). This study, however, fail to detect *DPLs* and *Sa*. Considering that *DPL* system is more likely an RI between *aus* and *temperate japonica*, we suggest parents for crossing experiment may affect gene mapping.

We found domestication had affected the genetic background of *DPLs* in rice. The pattern of π and Hd reversed after domestication: in the two wild ancestors, the diversities of *DPL1* was higher than *DPL*2 while in *O. sativa* we found the opposite. We believe the reason is domestication and during the process, the two loci *DPL1* and *DPL2* underwent different process.

After found the Hd of DPL2- reduced badly, we surprisingly found the frequency of *DPL2-* was much higher in *temperate japonica* (87.8%) than the ancestral population rGDGX (40.9%). Similarly, the frequency of *DPL1-* was 45.0% in nIND and then increase to 70.6% in *aus*, but the Hd of DPL1- reduced in *aus* (Fig. 4). The results proves that *DPL1-* and *DPL2-* somehow were preferred by artificial selection, even the haplotypes were randomly reserved and underwent diversity reduce.

Considering our results, we assume *DPL1* and *DPL*2 may share similar function hence one of the copies is redundant. Among populations the lack of polymorphic sites is commonly found at either *DPL1* or *DPL*2 (Table 1), but hardly occurs at both loci. Furthermore, we found in *aus*, which carries a high frequency of *DPL1-*, the *DPL*2 is highly conserve with lower π and Hd than other groups. The same pattern occurs on *temperate japonica* carrying the highest frequency of *DPL2-*, the *DPL1* of which show a reduce of diversity.

More evidences are still required to answer how the selection formed the *DPL* system. We failed to detect significant results of Tajima’s D in *O. sativa* while the lack of polymorphic sites was found at *DPL1* locus of *aromatic*, *rayada* and *temperate japonica*, indicating the selection pressure differed among groups. In the wild ancestors purifying selection was detected, however we considered the results were false caused by intraspecies structure. Significant negative values of Tajima’s D is a sign of extremely low polymorphism and considered as a consequence of purifying selection, but populations expansion or unknown structure can also lead to significant negative (Tajima, 1989). Therefore, the neutral tests are designed for one population with fluent gene flow and not suitable for the widespread species. As *O. rufipogon* and *O. nivara* have strong structure within and between species (Liu et al, 2015; Londo et al, 2006), false significant negative are likely to be detected. Similar phenomenon was found on *S5*: the neutral test results were also significant negative in one wild ancestor (*O. rufipogon*) but not in *O. sativa* (Du et al, 2011). We suggest the reason may be also interspecies structure of wild species and detailed study among the groups of *O. sativa* to clear the role of artificial selection.

### Ancestral structure is the main force behind the establishment of *DPL* system

The results of Phylogeography proved that the intraspecies structure of wild ancestors played an important role to shape *DPL* system, if not the crucial one. We found the H_1 of *DPL1* network contained wild rices mostly from *O. nivara* of India and *O. rufipogon* from Oceania. India is the possible domestication centre of *aus* (Civan et al, 2015; Khush, 1997; Vaughan et al, 2008) and closely connected with domestication of *O. sativa* (Kovach et al, 2007; Morishima et al, 1992; Sweeney and McCouch, 2007; Vaughan et al, 2008). The *O. rufipogon* haplotypes in H_13 of *DPL*2 network came from South China (rGDGZ and rGXHZ), where was proved as the origin centre of *Japonica* (Civan et al, 2015; Khush, 1997; Londo et al, 2006; Vaughan et al, 2008). We assumed the ancestral population of *DPL1-* was nIND and these for *DPL2-* were rGDGZ and rGXHZ.

Mizuta et al (2010) assumed *DPLs* were a barrier between the two subspecies, *Indica* and *Japonica*. However, the study did not separate *aus* from *indica* and their result actually indicated *DPL1-* clustered in *aus*, not in *indica*. The study of Craig et al (2014) and our results further proved that *DPL1-* took a high frequency in *aus* but rare in *indica*. Parsimoniously, we assume that *aus* inherited *DPL1-* directly from wild ancestors, then spread it among other groups. Our study proved that *DPL2-* had the highest frequency in *temperate japonica* (87.8%). The groups belonging to *Japonica* (*aromatic*, *rayada*, *temperate japonica* and *tropical japonica*) have higher frequencies than *Indica* groups (*aus* and *indica*) (Table 1). We hence assume the *Japonica* groups (highly probable *temperate japonica*) inherited *DPL2-* first, and then spread it into other groups. In conclusion, it needs at least twice domestications for *O. sativa* to obtain *DPL1-* and *DPL2-*. Hence our study is in accord with the “multiple domestications” hypothesis, by which *O. sativa* is believed as a result of more than one domestications (Sang and Ge, 2007; Yang et al, 2012a).

Similar to *DPL1-*, an *Indica-*like haplotype H_9 of another RI system, *S5*, had a discontinuous distribution: existed at New Guinea and South Asia, but not at anywhere in between (Du et al, 2011). Nevertheless, the study did not located any *Japonica-*like haplotype of *S5* system. The *Japonica* haplotype (*SaM-SaF-*) of *Sa* system was proved originate from an *O. rufipogon* population in southern China yet no evidence for the origination area of *Indica* haplotypes (Long et al, 2008). Our study for the first time confirms the original centres of both parts, and proves that the RI between *Indica* and *Japonica* were formed by ancestral geographic pattern.

Not only in rice, the effects of ancestral geographic pattern on RI genes are widely found. The same mechanism occurs in other *Oryza* species. *S27*/*S28* cause hybrid pollen sterility between *O. sativa* and *O. glumaepatula*. The system were under the influence of the ancestral geographic pattern and rapid sampling effects during the divergence of the two species (Yamagata et al, 2010). In the ancestor of two selfing descendants *Capsella rubella* and *C. grandiflora*, the balancing selection maintains the polymorphism of two potential incompatible loci *NPR1* and *RPP5*, facilitate the establishment of RI (Sicard et al, 2015).

### The *DPLs* are maintained in the wild ancestors because of the absence of selection and geographic isolation

RI genes lead to hybrid incompatibility, which means a certain number of descendants fail to survive or breed. Therefore, RI genes seem deleterious for the populations and are expected to loss, especially in the condition of gene flow (Bank et al, 2012; Gavrilets, 1997). On the contrary, RI loci are commonly found polymorphic, especially in plant (Bank et al, 2012; Corbett-Detig et al, 2013; Cutter, 2012; Lindtke and Buerkle, 2015; Rieseberg and Blackman, 2010). Here comes the question that how the incompatible alleles can be maintained in facing alternative alleles.

We found *DPL1-* and *DPL2-* were maintained because of geographically isolation (Table 1). Besides, neither of *DPL1-* nor *DPL2-* is deleterious along. In *O. glaberrima* and its wild ancestor *O. barthii*, *DPL1* totally pseudogenelizes because of losing the longest exon (Mizuta et al, 2010). Two *O. nivara* populations of Laos in our study, *DPL1-* takes high frequency (100% and 90.5%, respectively). *DPL2-* also takes 34.8% and 47.6% at two *O. rufipogon* populations in South China. So far, none of the species or populations shows recession.

The examples about ancestral polymorphism leading to RI systems are found in many species. Loci leading to male sterility are variance within grasshopper populations, proves that selection on incompatible loci could be weak, even absent(Shuker et al, 2005). Furthermore, the RI loci of *Capsella* species are proved under balance selection (Sicard et al, 2015). In the ancestor *C. grandiflora*, the balancing selection maintain the polymorphism of two potential incompatible loci *NPR1* and *RPP5*, facilitate the establishment of RI between the two selfing descendants, *C. rubella* and *C. grandiflora*. A pair of closely linked genes causes a globally distributed incompatibility in *Caenorhabditis elegans*, and is maintained by balance selection (Seidel et al, 2008). It is presumed that purifying selection may also be helpful for polymorphism. *Hl13* and *hl14* cause hybrid lethality between *Mimulus guttatus* and *M. nasutus* and both loci are polymorphic in the two species (Zuellig and Sweigart, 2018). The incompatibility alleles are maintained for decades. The authors assume that continual gene flow keep purging the lethal combinations and replenishing polymorphism at both loci.

Our result also implies that the establishment of *DPL* system has no concern with the gene function, but relies on the increase of frequency of non-functional alleles. In this hypothesis, the ancestral polymorphism is a crucial precondition. Other than the non-functional alleles detected by Mizuta et al (2010), we found other alleles that may possibly loss function. For example, a 7 bp insertion in the longest exon occurs only in wild rices from Sri Lanka. It is plausible that the RI caused by *DPLs* may work in a more complex way in the natural condition. We assume that if the frequency of any other non-functional alleles increases after domestication, the RI can still establish. We hence hypothesis that the RI systems of rice or even in other species, may arise in similar way as *DPLs*. We also suggest that analysis on population level is necessary to clarify the evolutionary history of RI systems, especially for widespread species.

## List of abbreviations

*DPL*: *DOPPELGANGER*
RI: reproductive isolation
IDN: Indonesia
IND: India
KHM: Cambodia
LAO: Laos
LKA: Sri Lanka
MMR: Myanmar
NEP: Nepal
PNG: New Guinea
THA: Thailand
GDGZ: Gaozhou, Guangdong
GXHZ: Hezhou, Guangxi
GXBH: Beihai, Guangxi
GXTD: Tiandong, Guangxi
HNDZ: Danzhou, Hainan
HNWC: Wenchang, Hainan
HNCL: Chaling, Hunan
HNJY: Jiangyong, Hunan
JXDX: Dongxiang, Jiangxi
YNJH: Jinghong, Yunnan
ARO: aromatic
AUS: aus
IND: indica
RAY: rayada
TEJ: temperate japonica
TRJ: tropical japonica
AMOVA: analysis of molecular variance
CDS: coding sequence
Hd: haplotype diversity
IRRI: the International Rice Research Institute

## Additional Files

**Additional File 1: Table S1.** Material list.

**Additional File 2: Table S2.** List of download *DPLs* sequence.

**Additional File 3: Figure S1.** The sketches of *DPLs* and the positions of primers.

## Declarations

### Ethics approval and consent to participate

Not applicable.

### Consent for publication

Not applicable.

### Data Archiving

All data generated or analysed during this study are included in this published article and its supplementary information files. All sequence data used in this article are available in GenBank (http://www.ncbi.nlm.nih.gov) under the accessions MN446022 - MN446138, MN446139 - MN446234, MN446235 - MN446531, MN446532 - MN446716, MN476103 - MN476405, MN476406 - MN476830, MN476831 - MN476868, MN476869 - MN476888.

### Authors’ contributions

F-MZ conceived and supervised the study. XX performed all experiments. XX and F-MZ analysed the data. F-MZ and XX wrote the manuscript with the help of GS. All authors have read and approved the final version of the manuscript.

## Acknowledgements

We are grateful to Cheng-Bin Chen from Guangxi academy of agricultural sciences for providing *O. rufipogon* populations of China. We are grateful to Xiao-ming Zheng from Beijing academy of agricultural sciences for providing *O. sativa* landraces. We are grateful to Xiao-ming Zheng, Hai-fei Zhou, Rong Liu, Lian Zhou, Qing-Lin Meng collected samples from natural habitats. We also thank the International Rice Research Institute (Los Banos, Philippines) for providing seed samples.

