## Supplementary figures and images for "Independent domestications shape the genetic pattern of a reproductive isolation system in rice"

### Additional file 3

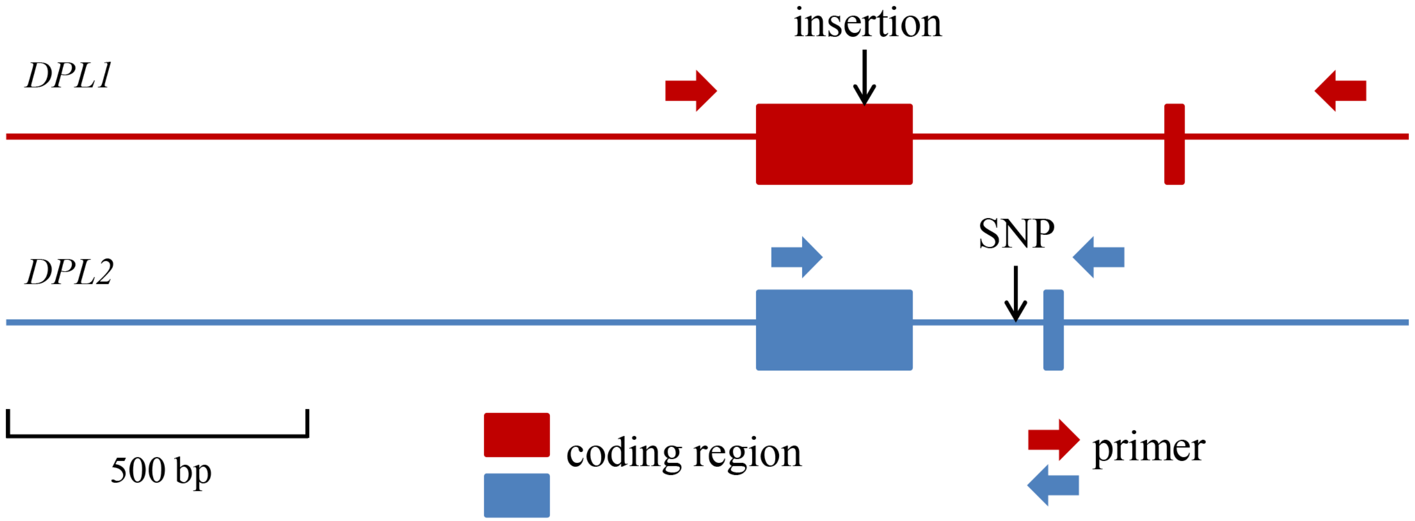
